# A comprehensive identification protocol for postcranial bones of Southeast Asian Ruminantia. Part 1: Osteometry

**DOI:** 10.64898/2025.12.18.695071

**Authors:** Papavee Seesod, Supalak Mithong, Sirikanya Chantasri, Corentin Bochaton

## Abstract

Taxonomic identification of animal bone remains from archaeological and paleontological contexts is usually conducted by comparison with modern skeletal specimens, an approach that is relatively straightforward in temperate or insular regions. In contrast, it is more challenging in continental tropical areas, where rich biodiversity makes it difficult to assemble exhaustive modern references. This situation is well illustrated in continental Southeast Asia, where studies of large mammal assemblages have largely focused on teeth and cranial elements, while postcranial remains are rarely investigated. To address this issue, we established an osteometric reference dataset of the limb bones from Southeast Asian ruminant species and analyzed both the potential of raw measurements and osteometric ratios to identify the different taxa. The results reveal marked differences between genera and species, allowing size-based grouping and precise taxonomic identification in some cases. These data were subsequently used to develop an easy-to-use identification tool based solely on measurements. The performance of this tool was tested against previously published datasets, allowing confirmation of existing results and discussion of potential biases associated with identification based on measurements alone. This new methodological framework provides a solid foundation to be combined with the qualitative morphological criteria described in the other papers of this series to enhance the precision and robustness of the taxonomic identifications of subfossil bones from continental Southeast Asia.

## Introduction

During the Late Quaternary, most vertebrate species were similar to their modern counterparts. Because of this, the identification of subfossil bone remains relies on direct comparison with reference skeletons of all species currently living in the biogeographic area where the remains were found. This approach makes research relatively straightforward in temperate or insular regions, which tend to have low vertebrate diversity. However, it becomes far more challenging in tropical regions, where species richness is much higher. Assembling a reference collection with several specimens per species—enough to document morphological variability—is manageable in areas with only a few dozen species, but it becomes far more difficult in regions that host several hundred and often lack major museum institutions. As a result of these constraints, most vertebrate groups remain poorly studied in the tropical Late Quaternary fossil record.

In continental Southeast Asia, for example, mammals have been the primary focus of most studies due to their abundance in the paleontological and archaeological records (Louys et al., 2007; Conrad, 2015). This emphasis is partly the result of the fact that much of the available data derives from the study of teeth alone (Bacon et al., 2008, 2011; Zeitoun et al., 2019; Suraprasit et al., 2021). Teeth are robust, often complete, relatively easy to study, and preserve well in most depositional contexts. However, this situation generates two major biases. First, taxa that lack large teeth are systematically underrepresented (Bochaton et al., 2024). Second, zooarchaeological assemblages—which are typically dominated by postcranial elements and can provide insight into the behaviors of past human populations—are only partially understood. These biases might be unavoidable if non-dental remains were genuinely rare in continental Southeast Asia, but this is not the case. The main obstacle to their study is the current lack of reference collections and dedicated research.

Although several studies have begun to disseminate reference data for the region’s mammal species, these efforts have mainly focused on cranial and dental morphology within paleontological and taxonomic frameworks (e. g. Heckeberg, 2020; Calamari, 2021). In contrast, reference works for postcranial elements and for non-mammalian taxa remain scarce. Our research group has begun addressing this gap for certain taxa, particularly turtles (Bochaton et al., 2023, 2025) and monitor lizards (Bochaton, Hanot, et al., 2019; Bochaton, Ivanov, et al., 2019). Nevertheless, even for the large mammals that dominate subfossil assemblages, substantial work is still needed to achieve adequate anatomical coverage and to enable reliable identification of remains in the absence of teeth.

To address this issue, our research team has developed a series of reference works focusing on the postcranial osteology of Ruminantia from continental Southeast Asia. This taxonomic group was chosen because ruminants constitute the majority of identifiable bone remains in Late Quaternary paleontological and archaeological deposits across the region. These studies aim to provide a set of methods that can be easily applied by researchers working without access to comprehensive skeletal reference collections. We also seek to make these methods as clear and reproducible as possible in order to improve the quality of regional zooarchaeological analyses, which are often based on limited comparative material of low representativeness. This paper is the first in a series of three. It focuses on the use of osteometric measurements for taxonomic identification, while the two subsequent papers address qualitative morphological characters (Chantasri et al., 2026; Mithong et al., 2026).

Osteometric studies of skulls and teeth from modern Southeast Asian ruminants have already been undertaken, either to support observations made on fossil specimens (Hooijer, 1958; Gruwier et al., 2015) or for taxonomic purposes (Guzmán & Rössner, 2018). However, only a few studies have examined postcranial elements, including measurements (Hooijer, 1958; Suraprasit et al., 2016). The most recent of these (Suraprasit et al., 2016) provides measurements and ratios for a substantial assemblage of modern Southeast Asian ruminants. Our study adopts a similar approach but uses a broader dataset—both in terms of the number of species and comparative specimens—while employing a more limited set of measurements.

The present study aims to provide an easy-to-use tool and a reproducible method that allows taxonomic attribution of bone elements based solely on measurements. To achieve this, we adopted a simple approach relying on 2D measurements that can be easily collected in practical research settings, and we developed an HTML-based tool that facilitates comparison between newly recorded measurements and the reference data presented here. We then applied this tool to three published datasets—two archaeological and one paleontological—to demonstrate its potential for improving the precision and reproducibility of taxonomic identifications in the study of subfossil bone remains.

## Material and Methods

### Selection of the investigated species and taxonomic remarks

The Ruminantia species included in this study were selected based on their documented occurrence in continental Southeast Asia (Lekagul & McNeely, 1988; Francis, 2019) or in recent paleontological studies (Suraprasit et al., 2016). All genera currently present in the region were sampled, except *Pseudoryx nghetinhensis* Dung, Giao, Chinh, Tuoc, Arctander & MacKinnon, 1993, for which no complete skeleton is available in museum collections. In a few cases, closely related species from outside the region were included to increase sample size at the generic level (e.g., *Capricornis crispus*, *Muntiacus reevesi*).

The species examined belong to four families (Bovidae Gray, 1821; Cervidae Goldfuss, 1820; Moschidae Gray, 1821; and Tragulidae Milne-Edwards, 1864) and seven tribes and subtribes (Hassanin et al., 2012). The considered species are the following: Bovidae: Bovini - *Bos gaurus* (Hamilton Smith, 1827); *Bos javanicus* d’Alton, 1823; *Bos sauveli* Urbain, 1937; *Bubalus bubalis* (Linnaeus, 1758); Caprina - *Budorcas taxicolor* Hodgson, 1850, *Pseudois nayaur* (Hodgson, 1833); Ovibovina - *Capricornis crispus* (Temminck, 1844), *Capricornis sumatraensis* (Bechstein, 1799), *Naemorhedus goral* (Hardwicke, 1825); Cervidae: Cervini Goldfuss, 1820 – *Axis axis* (Erxleben, 1777), *Axis porcinus* (Zimmermann, 1780), *Cervus nippon* Temminck, 1838, *Rucervus eldii* (McClelland, 1842), *Rucervus schomburgki* Blyth, 1863, *Rusa unicolor* (Kerr, 1792); Muntiacini - *Elaphodus cephalophus* Milne-Edwards, 1872, *Muntiacus muntjak* (Zimmermann, 1780), *Muntiacus reevesi* (Ogilby, 1839); Moschidae: *Moschus moschiferus* Linnaeus, 1758; Tragulidae: *Tragulus* sp. Brisson, 1762.

The necessary details regarding the species definitions and the taxonomy followed in this study are provided below.

*Bos gaurus* is currently divided into two subspecies: *B. g. gaurus* (Hamilton Smith, 1827) in India and Nepal, and *B. g. laosiensis* (Heude, 1901) in Southeast Asia (Groves & Grubb, 2011; Garrick & Ruvinsky, 2015). The domestic form is treated separately and recognized as *Bos frontalis* (Gentry et al., 2004). During our data collection, we examined all skeletal specimens attributed to *Bos gaurus*—likely representing both subspecies—but did not include specimens assigned to *Bos frontalis*.

Similarly, *Bos javanicus* is considered to comprise two subspecies: *B. j. birmanicus* in continental Southeast Asia and *B. j. javanicus* on Java and Borneo (Garrick & Ruvinsky, 2015), along with a domestic form. The specimens we documented may correspond to any of these three forms.

The systematics of water buffaloes have recently undergone major revisions (see literature review in Zhang et al., 2020). Following the taxonomic split between wild (*Bubalus arnee*) and domestic (*Bubalus bubalis*) forms (Gentry et al., 2004), genetic studies have shown that several domestic lineages diverged before domestication (Curaudeau et al., 2021). Two domestic species are now recognized: *Bubalus bubalis* (Linnaeus, 1758), the river buffalo domesticated in India, and *Bubalus kerabau* (Fitzinger, 1860), the swamp buffalo domesticated in Southeast Asia. These findings render the status of the wild buffalo *Bubalus arnee* uncertain, as it may represent multiple wild lineages within these domestic species. Consequently, it is difficult to determine with precision which species many older museum specimens correspond to. In this study, we treat all specimens labelled *Bubalus bubalis* as belonging to the broad domestic–wild complex currently recognized within *Bubalus*, excluding the small-bodied species (*B. depressicornis* Smith, 1827; *B. mindorensis* Heude, 1888; and *B. quarlesi* Ouwens, 1910).

Regarding serows and gorals, recent molecular studies (Mori et al., 2019; Li et al., 2020) have shown that *Naemorhedus griseus* (Milne-Edwards, 1871) is conspecific with *Naemorhedus goral* (Hardwicke, 1825), and that *Capricornis milneedwardsii* David, 1869 is a synonym of *Capricornis sumatraensis* (Bechstein, 1799) (Mori et al., 2019). Accordingly, the museum specimens we examined were attributed to these species. No skeletal specimens of *Capricornis rubidus* (David, 1869) or *Naemorhedus baileyi* Pocock, 1914—both valid species—were present in the collections visited.

The systematics and phylogeny of Cervidae have been the subject of numerous investigations based on molecular, morphological, and combined evidence approaches (see literature review in Heckeberg, 2020). As a result, the relationships among modern taxa are now considered relatively well resolved. The eight cervid species included in this study belong to two distinct tribes (Gilbert et al., 2006): Muntiacini Knottnerus-Meyer, 1907 (genera *Muntiacus* Rafinesque, 1815, and *Elaphodus* Milne-Edwards, 1872) and Cervini Goldfuss, 1820 (genera *Cervus* Linnaeus, 1758, *Rucervus* Hodgson, 1838, and *Rusa* C. H. Smith, 1827).

Regarding *Muntiacus*, three of the five species currently present in continental Southeast Asia (Francis, 2019) were described within the past 30 years: *M. gongshanensis* in 1990 (Shi-lai et al., 1990), *M. truongsonensis* in 1998 (Giao et al., 1998), and *M. putaoensis* in 1999 (Amato et al., 1999). In the historical collections we examined, specimens were generally assigned to *M. muntjak* for Southeast Asian material and *M. reevesi* for Chinese material. We chose to retain these attributions, while acknowledging that each may encompass multiple species; nonetheless, they still provide reliable information at the genus level.

Within the tribe Cervini, several genera are currently recognized (IUCN, 2025) and well established for the species considered here: *Axis* Smith, 1827 (species *axis* and *porcinus*), *Cervus* Linnaeus, 1758 (species *nippon*), *Rusa* Hamilton Smith, 1827 (species *unicolor*), and *Rucervus* Hodgson, 1838 (species *eldii* and *schomburgki*). Their distinction is justified based on antler morphology (Heckeberg, 2017) and, in some cases, dental characters (Meijaard & Groves, 2004a).

The family Moschidae includes only a single extant genus, *Moschus* (Groves & Grubb, 2011). This genus is represented by two species in continental Southeast Asia: *Moschus fuscus* Li, 1981 and *Moschus berezovskii* Flerov, 1929 (Francis, 2019). Modern classifications of the genus recognize a varying number of species (Zhang et al., 2026), but earlier classifications lumped most of these under a single name, *Moschus moschiferus* Linnaeus, 1758. Consequently, *M. moschiferus* was, in most cases, the only species available in the historical museum collections from which we defined our characters. As with *Muntiacus*, these specimens might correspond to several species of *Moschus*, and our observations should therefore be considered relevant only at the genus level, rather than as characters of *Moschus moschiferus* sensu stricto.

The taxonomy of *Tragulus* has undergone numerous revisions since the early 20th century. Initially, only two species were recognized: the lesser mouse-deer (*T. javanicus*) and the greater mouse-deer (*T. napu*) (Smit-van Dort, 1989). These “species” were later shown to comprise two distinct species groups distributed across continental and insular Southeast Asia (Meijaard & Groves, 2004b). In the museum collections examined for this study, nearly all specimens were labelled either *T. javanicus sensu lato* or *T. napu sensu lato*. To avoid taxonomic uncertainty, we therefore chose not to treat these forms separately and restricted our observations to the genus level for *Tragulus*.

A total of 286 specimens representing 20 taxa were measured and included in this study. For each specimen, we recorded as many measurements as possible while avoiding damaged areas and bones with remaining dried soft tissues. The species selected for measurement were chosen based on their modern or historical occurrence in Thailand and the surrounding regions (Lekagul & McNeely, 1988). Our datasets include the following taxa: *Axis axis* (N=11); *Axis porcinus* (N=8); *Bos gaurus* (N=16); *Bos javanicus* (N=14); *Bos sauveli* (N=3); *Bubalus bubalis* (N=21); *Budorcas taxicolor* (N=18); *Capricornis crispus* (N=7); *Capricornis sumatraensis* (N=12); *Cervus nippon* (N=18); *Elaphodus cephalophus* (N=11); *Moschus moschiferus* (N=11); *Muntiacus muntjak* (N=24); *Muntiacus reevesi* (N=14); *Naemorhedus goral* (N=12); *Pseudois nayaur* (N=20); *Rucervus eldii* (N=15); *Rucervus schomburgki* (N=1); *Rusa unicolor* (N=25); and *Tragulus* sp. (N=25). These specimens were obtained from five museum collections in France, Belgium, Germany, and the United States (see full details in Supplementary Material 1; Seesod et al., 2026).

### Measurements and analyses

A set of 59 measurements distributed across all girdle and limb bones was selected from the parameters commonly used in osteometric studies (Driesch, 1976) and recorded for our sample. These measurements were selected to provide a rough estimate of bone size and geometry. We deliberately limited the number of measurements recorded in order to measure as many specimens as possible in the visited collections. Because this study was conducted in parallel with the search for qualitative identification criteria on the same skeletal elements, bones of the axial skeleton were not included. The measurements recorded were as follows: Scapula (HS, Ld, SLC, GLP, BG); Humerus (GLC, Bp, SD, Bd); Ulna (LO, SDO); Radius (GL, Bp, SD, Bd); Scaphoid (GB); Lunatum (GB); Pyramidal (GB); Pisiform (GB); Magnum (GB); Unciform (GB); Metacarpus (GL, Bp, SD, Bd); Coxal (GL, LA); Femur (GLC, Bp, SD, Bd); Patella (GL, GB); Tibia (GL, Bp, SD, Bd); Malleolar bone (GD); Astragalus (GLI, DI, Bd); Calcaneus (GL, GB); Navicular-cuboid (GB); Large cuneiform (GB); Metatarsus (GL, Bp, SD, Bd); First phalanx (GL, Bp, Bd); Second phalanx (GL, Bp, Bd); Third phalanx (DLS, MBS); Small sesamoid (GB); and Large sesamoid (GB). These measurements are illustrated in Supplementary Material 2 (Seesod et al., 2026). Only phalanges and sesamoids were measured several times in some specimens to account for their intra-individual variability. All measurements were taken with a digital dial caliper [IP67 (Mitutoyo Corporation, Japan)] with a 0.01 mm measurement precision by all authors of this study.

After checking the dataset for measurement errors, the final data (Supplementary Materials 3 and 4; Seesod et al., 2026) were subjected to several statistical analyses and visualizations using R v. 4.6.0 (R Core Team, 2026) (https://cran.r-project.org), with the packages stats, tidyverse (Wickham & RStudio, 2023), and ggplot2 (Wickham et al., 2025).

To evaluate the specific and generic discriminatory power of both raw measurements and ratios, several variables were calculated for each measurement within each species and genus: the number of taxa (species or genera) whose measurement ranges overlap with that of the focal taxon (NTO), the total amount of overlap between the focal taxon and all other taxa (SO), the mean proportion of the focal taxon’s range that overlaps with each taxon showing non-zero overlap (MPO), and the proportion of the focal range falling outside the combined range of all other taxa (PRU). For the synthesis of character discriminatory power, we also calculated the mean overlap length across all taxa that overlap with the focal taxon (MOL). To provide an indication of relative size differences among taxa, we computed the mean relative range position (RRP) for each measurement, which shows whether a taxon tends to fall toward the lower or upper end of the overall measurement spectrum. These calculations were performed using an R script (Supplementary Material 5; Seesod et al., 2026) initially generated using Claude AI (Anthropic Opus 5 Model) and manually checked and revised afterward. For variables represented by a single available value, a pseudo-range of ±0.005 mm was assigned to avoid disrupting the calculations.

Finally, we performed a Principal Component Analysis on the length of each long bone (humerus, radius, femur, tibia, and metapodials) as well as the scapula, to investigate proportional differences among the studied taxa. The raw data were first transformed to Mosimann shape variables (Mosimann & James, 1979), calculated from natural log-transformed measurements. All of these analyses were conducted using the provided R script (Supplementary Material 5; Seesod et al., 2026).

To conclude our study and to test the applicability of our measurement and ratio references for identifying paleontological and archaeological remains, we developed an easy-to-use HTML tool (Bochaton et al., 2026) using Claude AI (Anthropic Opus 5 Model), then manually checked and revised afterward. After the user inputs a set of measurements and/or ratios, the tool automatically identifies the taxa that match these values and indicates the most likely attribution based on our dataset. To do so, the tool ranks the taxa by decreasing proportion of compatible values, those compatible with every comparable value being listed first, ties being broken in favour of the taxa evaluated on more variables and documented by more reference specimens.

We then used this tool to evaluate previously published identifications based on bone measurements and ratios. For this purpose, we selected two zooarchaeological studies (Grant & Higham, 1991; Auetrakulvit, 2004) and one Quaternary paleontological study (Suraprasit et al., 2016), all of which provide raw measurements of bones attributed to Ruminantia. For each bone, we retained the identification(s) matching the input data at 100%, those exceeding 80%, or, when no higher matches were available, those with lower matching percentages. Only the best or equally best matching identifications were considered.

## Results

The following results were obtained using the R script provided in Supplementary Material 5 (Seesod et al., 2026). They are based on the measurement ranges (Supplementary Materials 6 and 7; Seesod et al., 2026) and ratios (Supplementary Materials 8 and 9; Seesod et al., 2026) for both genera and species, as well as on their corresponding overlap values (Supplementary Materials 10–13; Seesod et al., 2026). The PCA was performed on the subset of specimens for which the lengths of all long bones were available (Supplementary Material 14; Seesod et al., 2026).

### Analysis of the raw measurements

At the generic scale (Table 01), the raw measurements show a substantial mean NTO value (4.64), indicating that each taxon generally overlaps with about one-third of all genera included in the sample. The overall MPO is also high (52%). For nearly all genera, measurement ranges overlap extensively, with mean PRU values equal to or below 1%. The exceptions are *Tragulus*, which almost never overlaps with other genera (mean PRU = 97%) due to its very small size (mean RRP = 3%), and *Bos*, whose larger values produce a relatively high mean PRU (28%). The three largest genera—*Bos*, *Bubalus*, and *Budorcas* (mean RRP = 63–77%)—rarely overlap with smaller taxa and consequently display a low mean NTO (3). All other genera overlap with at least 3.5 others on average, except *Moschus*, which shows limited size variability in our sample. Despite this overlap, the mean RRP values reveal clear size differences among groups. In addition to *Tragulus* and the three large-bodied genera mentioned above, a size gap separates the smaller remaining genera (*Axis*, *Elaphodus*, *Moschus*, *Muntiacus*, and *Naemorhedus*, mean RRP = 16–29%) from the larger ones (*Capricornis*, *Cervus*, *Rucervus*, and *Pseudois*, mean RRP = 31–38%). *Rusa* (mean RRP = 48%) is in an intermediate position between the latter and the large bovids group. Based on these patterns, which vary slightly among individual measurements, five size classes can be distinguished (Figure 1) with *Rusa* and *Tragulus* forming an intermediate and an extreme size group on their own.

**Figure 1.**
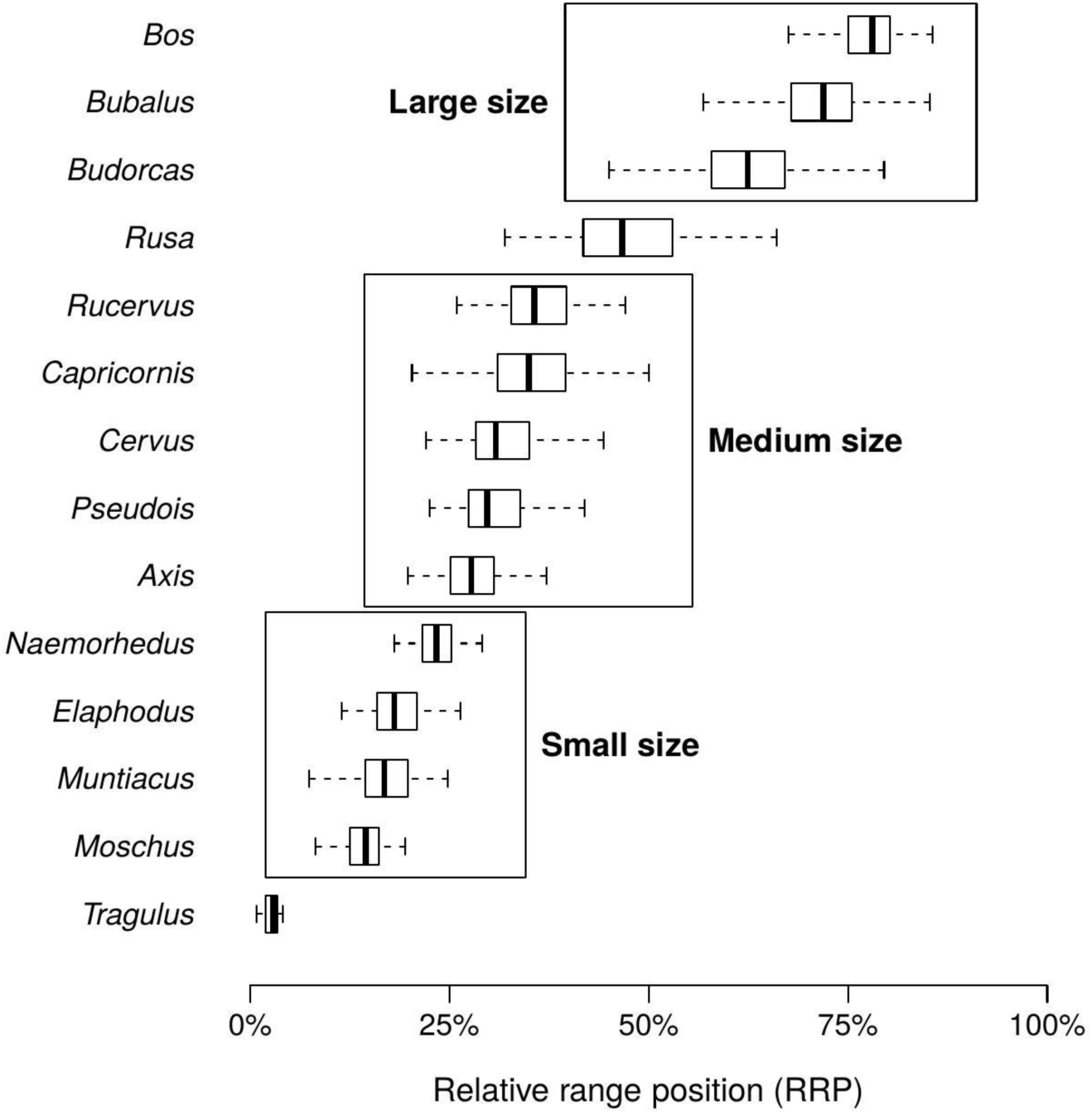
Plot of the relative range position (RRP) values of the different measurements for each taxon showing size differences between them. Data from Supplementary Material 10 (Seesod et al., 2026).

**Table 01.** Discriminatory potential of the raw measurements among the different genera.

| Genus | NTO | SO | MOL | MPO | PRU | RRP |
| --- | --- | --- | --- | --- | --- | --- |
| *Axis* | 7.14 | 4021.06 | 9.27 | 51% | 0% | 29% |
| *Bos* | 2.92 | 3341.21 | 19.46 | 41% | 28% | 77% |
| *Bubalus* | 2.98 | 3280.5 | 17.77 | 62% | 1% | 72% |
| *Budorcas* | 3.25 | 2859.76 | 14.29 | 59% | 1% | 63% |
| *Capricornis* | 6.86 | 4535.38 | 11.04 | 47% | 0% | 36% |
| *Cervus* | 7.83 | 5481.3 | 11.58 | 43% | 0% | 32% |
| *Elaphodus* | 3.6 | 619.62 | 2.96 | 67% | 0% | 19% |
| *Moschus* | 2.31 | 651.4 | 4.16 | 78% | 1% | 16% |
| *Muntiacus* | 5.95 | 2006.93 | 5.51 | 27% | 1% | 17% |
| *Naemorhedus* | 5.27 | 1630.68 | 5.09 | 65% | 0% | 24% |
| *Pseudois* | 5.73 | 3108.6 | 9.51 | 69% | 0% | 31% |
| *Rucervus* | 5.73 | 3682.99 | 11.37 | 59% | 0% | 38% |
| *Rusa* | 5.17 | 2706.05 | 8.38 | 37% | 0% | 48% |
| *Tragulus* | 0.18 | 17.09 | 1.12 | 18% | 97% | 3% |
| Mean | 4.64 | 2710.18 | 9.39 | 52% | 9% | / |

The comparison among the different bones does not reveal many meaningful variations (Supplementary Material 15; Seesod et al., 2026). The mean NTO ranges from 2.17 to 9.86, but generally falls between 4 and 5. The other variables are mostly influenced by biases such as overall bone size, which affects SO and MOL, or the presence/absence of measurements for *Tragulus*, which strongly impacts the mean PRU (e.g., for the large sesamoids). Nonetheless, the third phalanx shows greater size variation among taxa than most other elements (mean NTO = 3.14). A similar pattern is observed for the large (proximal) sesamoids, although these were not measured for all species (only 14/20) and not for *Cervus* and *Tragulus*. Overall, however, all bone elements produce broadly similar results.

At the species level (Table 02), the raw measurement values show a substantial mean NTO (5.14), indicating an average overlap with about one quarter of the species (N = 20) included in the dataset. The MPO is high (55%). The mean PRU is below 1% for most species, except for *Tragulus* sp. (97%) and *Bos gaurus* (31%), which are respectively the smallest and largest species in the dataset. All other species exhibit strong size overlaps, with a mean of 5.4 overlapping species per measurement, each overlap representing, on average, 55% of the focal species’ measurement range. This confirms that most species share considerable size overlap, although size classes remain distinguishable. Large species show clear differences in mean RRP—*Bos gaurus* (81%), *Bubalus bubalis* (72%), *Bos javanicus* (69%), *Bos sauveli* (63%), and *Budorcas taxicolor* (63%)—indicating distinct size categories despite extensive overlaps that prevent reliable discrimination in many cases. *Rusa unicolor* (mean RRP = 48%) occupies an intermediate position with *Rucervus schomburgki* (46%), smaller than the large bovids but larger than the remaining taxa. Below it are *Capricornis sumatraensis* and *Rucervus eldii* (mean RRP = 39% and 36%), followed by *C. nippon*, *Axis axis*, *Pseudois nayaur*, and *Capricornis crispus* (mean RRP = 32–28%). Next are *Naemorhedus goral* and *Axis porcinus* (mean RRP = 24–23%), then *Elaphodus cephalophus*, *Muntiacus muntjak*, and *Moschus moschiferus* (mean RRP = 19–16%). *Muntiacus reevesi* (mean RRP = 12%) is the smallest taxon after *Tragulus* sp. (mean RRP = 3%). Across anatomical elements, the observed patterns are broadly consistent with those identified at the generic scale (Supplementary Material 16; Seesod et al., 2026).

**Table 02.** Discriminatory potential of the raw measurements among the different species.

| Species | NTO | SO | MOL | MPO | PRU | RRP |
| --- | --- | --- | --- | --- | --- | --- |
| *Axis axis* | 7.1 | 2828.75 | 6.81 | 54% | 0% | 32% |
| *Axis porcinus* | 5.31 | 1114.88 | 3.46 | 67% | 0% | 23% |
| *Bos gaurus* | 3.73 | 3839.17 | 16.67 | 42% | 31% | 81% |
| *Bos javanicus* | 4.75 | 4055.33 | 14.84 | 56% | 0% | 69% |
| *Bos sauveli* | 3.81 | 1555.57 | 6.73 | 80% | 0% | 63% |
| *Bubalus bubalis* | 4.86 | 4546.38 | 15.11 | 52% | 1% | 72% |
| *Budorcas taxicolor* | 4.9 | 3710.19 | 12 | 49% | 1% | 63% |
| *Capricornis crispus* | 5.73 | 1142.97 | 3.41 | 72% | 0% | 28% |
| *Capricornis sumatraensis* | 6.42 | 2758.1 | 7.42 | 49% | 0% | 39% |
| *Cervus nippon* | 10.17 | 5325.75 | 8.62 | 32% | 0% | 32% |
| *Elaphodus cephalophus* | 4.03 | 646.63 | 2.85 | 64% | 0% | 19% |
| *Moschus moschiferus* | 3.14 | 772.03 | 3.8 | 70% | 1% | 16% |
| *Muntiacus muntjak* | 7.55 | 2370.41 | 5.06 | 27% | 0% | 18% |
| *Muntiacus reevesi* | 2.28 | 530.38 | 5.1 | 68% | 0% | 12% |
| *Naemoredus goral* | 6.05 | 1661.06 | 4.49 | 58% | 0% | 24% |
| *Pseudois nayaur* | 7.07 | 2904.81 | 7.56 | 54% | 0% | 31% |
| *Rucervus eldii* | 6.27 | 3205.07 | 8.93 | 57% | 0% | 36% |
| *Rucervus schomburgki* | 2.7 | 1.43 | 0.01 | 100% | NA | 46% |
| *Rusa unicolor* | 6.64 | 2777.08 | 6.28 | 28% | 0% | 48% |
| *Tragulus sp.* | 0.21 | 24.43 | 1.23 | 18% | 97% | 3% |
| **Mean** | **5.14** | **2288.52** | **7.02** | **55%** | **7%** | **/** |

### Ratio between two measurements

At the generic scale (Table 03), the ratio values show a very high mean NTO (9.97), indicating that each genus overlaps, on average, with more than two-thirds of the genera (N = 14) included in the dataset. The MPO is also comparatively high (61%). Unique values are nearly absent (mean PRU = 4%), although *Budorcas* (13%), *Moschus* (15%), and *Tragulus* (8%) present a significant proportion of values that are exclusive to their genera.

**Table 03.** Discriminatory potential of the ratio between two measurements for the different genera.

| Genus | NTO | SO | MOL | MPO | PRU |
| --- | --- | --- | --- | --- | --- |
| *Axis* | 11.02 | 301.25 | 0.49 | 61% | 1% |
| *Bos* | 9.07 | 192.13 | 0.4 | 48% | 5% |
| *Bubalus* | 7.88 | 139.71 | 0.41 | 62% | 6% |
| *Budorcas* | 8.38 | 142.22 | 0.28 | 64% | 13% |
| *Capricornis* | 10.85 | 260.03 | 0.41 | 60% | 1% |
| *Cervus* | 10.77 | 290.35 | 0.49 | 62% | 1% |
| *Elaphodus* | 10 | 219.53 | 0.4 | 69% | 0% |
| *Moschus* | 8.22 | 156.41 | 0.38 | 60% | 15% |
| *Muntiacus* | 11.33 | 351.64 | 0.56 | 56% | 1% |
| *Naemorhedus* | 10.25 | 201.18 | 0.34 | 69% | 0% |
| *Pseudois* | 10.55 | 229.92 | 0.39 | 66% | 0% |
| *Rucervus* | 10.27 | 255.59 | 0.45 | 64% | 2% |
| *Rusa* | 10.95 | 286.51 | 0.46 | 62% | 0% |
| *Tragulus* | 10 | 289.95 | 0.52 | 49% | 8% |
| **Mean** | **9.97** | **236.89** | **0.43** | **61%** | **4%** |

Some anatomical elements show differences in their discriminatory potential (Table 04). The patella performs best, with an NTO of 6.57 and a mean PRU of 9%. The metacarpus also has a relatively high mean PRU (7%), although its mean NTO is higher (7.98). In contrast, most other elements perform similarly, except for the astragalus and calcaneus, which show high mean NTO values (>12) and very low mean PRU (≤1%), indicating almost no differences in the proportions of these bones among genera. A closer examination of individual ratio–genus combinations (Supplementary Material 12; Seesod et al., 2026) reveals that some ratios present unique values for more than 50% of their range (N = 19), and in a few cases for 100% of their range (N = 8). These highly discriminating ratios occur only in *Tragulus* (patella), *Moschus* (phalanx 1, phalanx 2, and metatarsus), and *Budorcas* (metacarpus and metatarsus).

**Table 04.** Generic discriminatory potential of the different anatomical elements with the ratio between two measurements.

| Bone | NTO | SO | MOL | MPO | PRU |
| --- | --- | --- | --- | --- | --- |
| Calcaneus | 13 | 104.09 | 0.57 | 75% | 0% |
| Coxal | 11.14 | 117.89 | 0.75 | 55% | 3% |
| Femur | 10.1 | 293.56 | 0.37 | 58% | 3% |
| Humerus | 9.71 | 285.67 | 0.46 | 65% | 2% |
| Metacarpus | 7.98 | 230.82 | 0.42 | 55% | 7% |
| Metatarsus | 8.33 | 319.08 | 0.52 | 56% | 8% |
| Patella | 6.57 | 16.52 | 0.18 | 56% | 9% |
| First phalanx | 9.67 | 113.51 | 0.29 | 62% | 4% |
| Second phalanx | 7.43 | 52.01 | 0.18 | 61% | 5% |
| Third phalanx | 9.14 | 71.79 | 0.51 | 63% | 2% |
| Radius | 10.43 | 380.58 | 0.47 | 60% | 2% |
| Scapula | 11.91 | 892.31 | 0.52 | 62% | 3% |
| Astragalus | 12.62 | 82.32 | 0.15 | 68% | 1% |
| Tibia | 9.83 | 312.12 | 0.47 | 61% | 4% |
| Ulna | 11.57 | 44.17 | 0.26 | 62% | 3% |
| **Mean** | **9.96** | **221.1** | **0.41** | **61%** | **4%** |

At the species level (Table 05), the ratio values show a very high mean NTO (13.17), indicating that each species overlaps, on average, with about two-thirds of the species (N = 20) in the dataset. The MPO is also very high (59%). The mean PRU is only 3%, and the specific patterns mirror those observed at the generic scale.

**Table 05.** Discriminatory potential of the ratio between two measurements for the different species.

| Species | NTO | SO | MOL | MPO | PRU | RRP |
| --- | --- | --- | --- | --- | --- | --- |
| *Axis axis* | 14.63 | 278.55 | 0.34 | 58% | 1% | 46% |
| *Axis porcinus* | 14.38 | 246 | 0.31 | 65% | 0% | 46% |
| *Bos gaurus* | 11.97 | 208.24 | 0.35 | 50% | 3% | 33% |
| *Bos javanicus* | 12.35 | 198.37 | 0.31 | 54% | 1% | 34% |
| *Bos sauveli* | 9.96 | 106.83 | 0.21 | 70% | 2% | 35% |
| *Bubalus bubalis* | 11.42 | 181.21 | 0.33 | 52% | 6% | 32% |
| *Budorcas taxicolor* | 11.35 | 170.6 | 0.25 | 58% | 13% | 40% |
| *Capricornis crispus* | 12.5 | 173.54 | 0.25 | 68% | 1% | 53% |
| *Capricornis sumatraensis* | 13.62 | 196.83 | 0.26 | 63% | 1% | 45% |
| *Cervus nippon* | 15.13 | 342.48 | 0.41 | 54% | 1% | 49% |
| *Elaphodus cephalophus* | 13.55 | 261.68 | 0.36 | 62% | 0% | 55% |
| *Moschus moschiferus* | 10.87 | 179.33 | 0.35 | 54% | 15% | 63% |
| *Muntiacus muntjak* | 15.87 | 397.24 | 0.45 | 51% | 1% | 50% |
| *Muntiacus reevesi* | 14.75 | 318.66 | 0.39 | 59% | 0% | 51% |
| *Naemorhedus goral* | 13.88 | 243.75 | 0.31 | 62% | 0% | 50% |
| *Pseudois nayaur* | 14.32 | 272.56 | 0.34 | 58% | 0% | 48% |
| *Rucervus eldii* | 14.07 | 279.71 | 0.37 | 58% | 2% | 48% |
| *Rucervus schomburgki* | 9.6 | 5.26 | 0.01 | 96% | NA | 45% |
| *Rusa unicolor* | 15.5 | 341.31 | 0.39 | 54% | 0% | 45% |
| *Tragulus sp.* | 13.6 | 340.23 | 0.46 | 43% | 8% | 58% |
| **Mean** | **13.17** | **237.12** | **0.32** | **59%** | **3%** | **/** |

Differences among anatomical elements follow the same trends as at the generic level (Supplementary Material 17; Seesod et al., 2026), with generally higher mean NTO values and lower mean PRU values across all bone elements. This applies to both the better-performing parts—the patella (mean NTO = 10.3; mean PRU = 6.9%) and the metacarpus (mean NTO = 10.80; mean PRU = 5%)—and the poorer-performing parts, such as the astragalus (mean NTO = 15.73; mean PRU = 0.6%) and the calcaneus (mean NTO = 17.3; mean PRU = 0.5%). The details of the best-performing ratios at the species level are not discussed here, as they correspond to the same genera identified at the generic scale, and these genera are each represented by a single species in the dataset.

### Proportion differences between the different long bones

The PCA performed on the greatest lengths of seven bones from the girdle (scapula) and limbs (humerus, radius, metacarpus, femur, tibia, and metatarsus) (Figure 2) shows that PC1 is dominated by the relative length of the metapodials. Metapodials are shortest in *Budorcas*, longer in other Bovidae, and even more elongated in Cervidae and Moschidae. PC2 appears to carry strong size information, as most taxa are arranged along this axis according to overall body size. *Budorcas* is clearly separated from the other taxa in the PC1–PC2 space due to its combination of elongated stylopods and zeugopods and shortened metapodials. Most cervids follow a common allometric trajectory. Moschidae differ by having proportionally longer metapodials than small cervids, while among larger-bodied taxa, Bovidae display relatively shorter metapodials than cervids of comparable size. Overall, the PCA indicates that, except for *Axis*, most genera are relatively well separated and show clear proportional differences among limb segments. Although these patterns are not independent of size—being influenced by allometry—this multivariate approach performs substantially better than analyses conducted on single bone elements.

**Figure 2.**
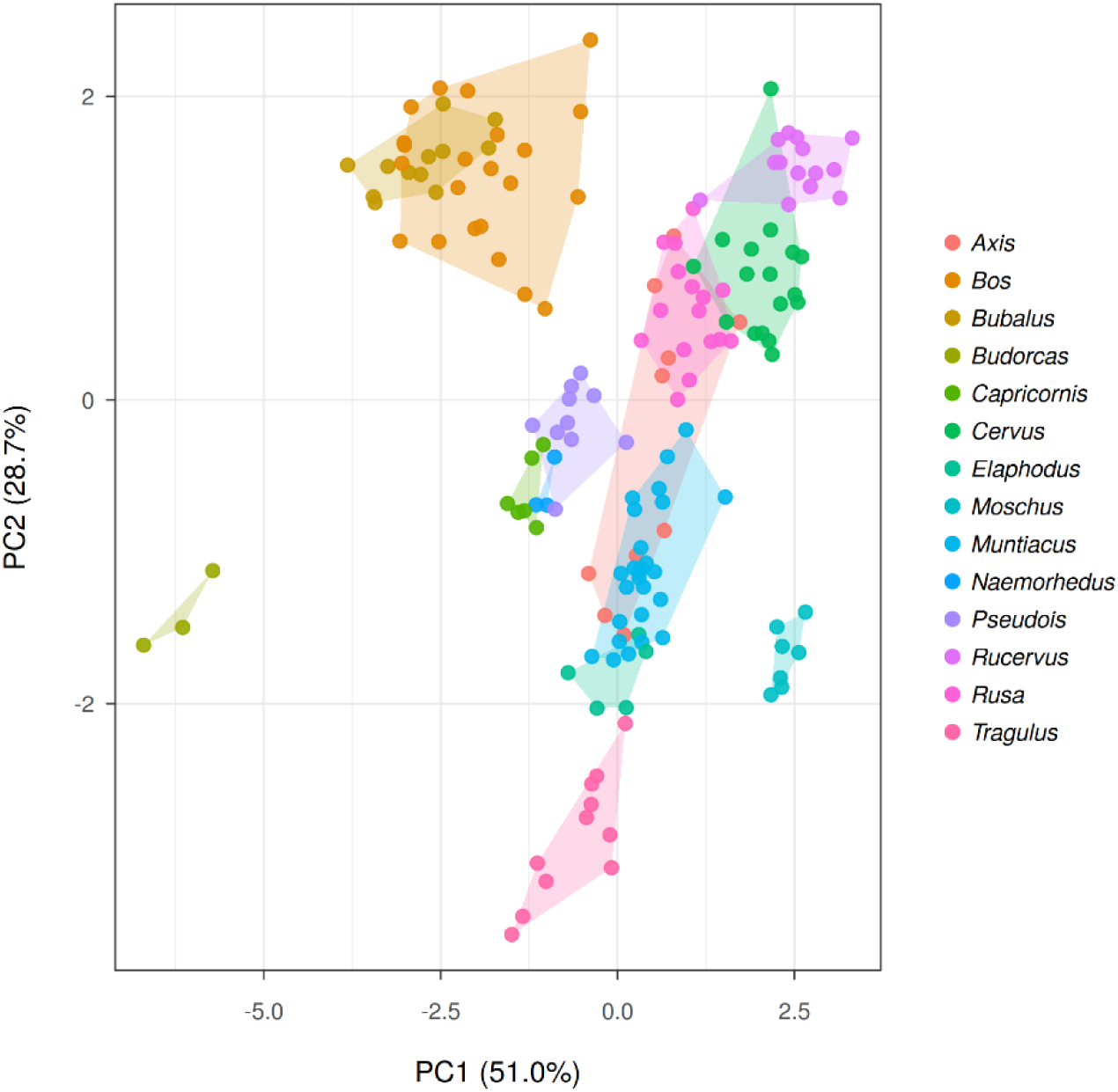
First two axes of the PCA of the scapula and the long bones’ greatest lengths.

### Application of our approach to published datasets

Applying our measurement and ratio baselines to existing datasets allowed us to identify 137 bones published in Auetrakulvit (2004), mostly phalanges, 95 bones from Grant and Higham (1991), and 56 bones from Suraprasit et al. (2016), mostly humeri, femora, metacarpus, and metatarsus. The results of these analyses are presented in Supplementary Material 18 (Seesod et al., 2026). On average, our method returned 2.6 equally most likely taxonomic identifications per bone, with at least 80% of variables matching in the vast majority of cases (247/288). This mean is lower (1.8) for the Suraprasit et al. (2016) dataset because more variables were available for each bone, and higher (3 and 2.4) for the other datasets because only a few measurements were available for each specimen. We tried to consider the raw measurements and the ratios separately, but this only results in a decrease in identification performance, especially in the case of the ratio, with a mean of 6 possible identifications per bone.

For the Auetrakulvit (2004) dataset, 15 bones provided inconclusive results (less than 80% of matching variables with a given taxon). The metric data yielded unambiguous identifications (a single possible taxon, matching 100% of the variables) in 28 cases. These include 21 bones previously identified at the family level only, of which 15 gave a result consistent with the original identification, 6 did not, and 7 bones previously identified at the genus or species level, of which 5 were consistent and 2 were not. For the remaining bones, our approach yielded 1 (N=3), 2 (N = 34), 3 (N = 17), 4 (N = 17), 5 (N = 13), 6 (N = 9), or 7 (N=1) fully matching possible attributions. Among the conclusive elements are 58 bones originally identified only at the family rank, for which our method provides greater taxonomic precision. The remaining cases (N = 36) correspond to bones previously attributed to genus or species; in these instances, our identification either agrees (N=2), agrees while indicating additional possible taxa (N = 22), or disagrees (N = 12). Overall, for this archaeological dataset, our results are largely congruent with the original interpretations, with only 16.4% discrepancies. In most cases, our method confirms or refines earlier identifications.

For the Grant and Higham (1991) dataset, only 13 bones provided inconclusive results (less than 80% of matching variables with a given taxon). The metric data yielded unambiguous identifications (a single possible taxon, matching 100% of the variables) in 21 cases. These are all bones previously identified at the genus or species level, of which 13 were consistent with our results, and 8 were not. For the remaining bones, our approach yielded 1 (N=4), 2 (N = 19), 3 (N = 17), 4 (N = 15), 5 (N = 5), or 6 (N = 1) fully matching possible attributions. Of these bones (N = 61), 60 had previously been attributed to genus or species and one to family only; in these instances, our identification either agrees (N=3), agrees while indicating additional possible taxa (N = 31), or disagrees (N = 27). Overall, for this archaeological dataset, our results are largely incongruent with the original interpretations, with 43% discrepancies. This is largely due to the high taxonomic precision of the original identifications, which makes them more likely to be contradicted than those in the previous dataset, but also to the fact that many of the bones attributed to *Rucervus eldii* present a size more matching that of *Rusa unicolor*.

Applying our approach to the Suraprasit et al. (2016) dataset proved more challenging because, although we use a wider modern comparative sample, we rely on fewer variables. In addition, the Middle Pleistocene age of the material increases the likelihood of morphological or size differences between modern specimens and their fossil counterparts. Finally, all bones in their study were identified to the species level, a resolution that our method generally cannot achieve. Despite these limitations, our approach provided unambiguous identifications (N possible taxa = 1 and 100% matching) for 21 bones. Six of these were originally identified as *Axis axis*; our results confirmed this attribution for two specimens but reassigned the remaining four to *Cervus nippon* and *Rusa unicolor*. Among the remaining 15 bones, our identifications match the original ones in 6 cases. The non-matching identifications correspond to 5 large bones originally attributed to *Bubalus* that our method identifies as *Bos gaurus*, 3 *Panolia eldii* (= *Rucervus eldii*) bones we attribute to *Cervus nippon*, and 1 *Rusa unicolor* bone we attribute to *Pseudois nayaur* (DMR-KS-05-04-19-10). For the remaining bones, our approach indicates 1 (N = 5), 2 (N = 6), 3 (N = 4), 4 (N = 2), or 5 (N = 5) significantly matching possible taxa. These include bones previously assigned to species for which our identifications either agree (N = 3), agree but introduce additional possible taxa (N = 12), or disagree (N = 7). In the full dataset, 13 bones did not provide identifications significantly matching any of our modern taxa. Among these are 8 bones originally attributed to *Bubalus arnee* that only partially match this species in our dataset. Overall, our results are largely incongruent with the original identifications, with 46% mismatches, but also introduce additional identification possibilities for 28% of the attributed elements. This is largely due to the high taxonomic precision of the original identifications, which makes them more likely to be contradicted than those in the previous datasets. Consequently, our conclusions are less definitive overall, and we were not able to confirm a substantial proportion of the identifications attributed to *Bubalus*.

## Discussion

### Comparison of the discriminatory performance of raw measurements and ratios

A general comparison between raw measurements and ratios shows that ratios perform poorly in most cases. This is clearly reflected in the mean NTO, which is more than twice as high for ratios than for raw measurements, indicating that although all variables show substantial overlap, the overlap is far greater for ratios. Ratios outperform raw measurements only in a few very specific cases, involving rare bones or particular taxa, but overall they underperform to such an extent that in some instances they are almost uninformative. Ratios related to bone gracility (i.e., combining length and breadth) do appear useful for distinguishing large bovids from cervids, although in our dataset they do not outperform single measurements. Beyond this, no ratio provides clearer taxonomic resolution than isolated measurements. We also note that ratio values for *Bos* and *Bubalus* overlap extensively, indicating that ratios are generally not effective for separating their postcranial remains, even though a few ratios perform relatively well. The most performing ratios are found on the metapodials, with the metacarpus showing more values without overlaps between the two genera (Bd/Bp – 93.2%; GL/Bd – 63.8%) than the metatarsus (Bd/Bp – 70.9%; GL/Bd – 55.4%; Bd/SD – 55.7%). All of the other ratios show less than half of their values not covered by the other genus. Although it has been suggested that the postcranial bones of *Bubalus arnee* are more massive and robust than those of *Bos* (Suraprasit et al., 2016), our results show that any such tendency is insufficiently pronounced to allow reliable identification based solely on gracility indices outside of the metacarpus and, to a lesser extent, the metatarsus.

### What our approach brings…

Our results show that identifying Ruminantia species from continental Southeast Asia using a simple 2D morphometric approach is feasible to some extent, and that these data can be informative even when not combined with qualitative morphological characters. The value of the approach we propose is illustrated by its application to published datasets: we were able to refine, confirm, or question existing identifications through a method that is both repeatable and open to scrutiny. This already represents progress, as size-based identifications in regional zooarchaeological studies are rarely supported by an adequate reference sample—unlike in most paleontological research. With this work, we formalize an approach that has been used informally for decades, providing a structured framework that will benefit scholars working in continental Southeast Asia, especially those lacking access to comprehensive modern skeletal reference collections, which are often absent locally. We also provide an easy-to-use tool (Bochaton et al., 2026) to facilitate the application of our method.

Regarding the applications shown here, our approach did not substantially modify the results of Auetrakulvit (2004); instead, we primarily added precision to existing identifications. However, several noteworthy observations emerge from the comparison with the Suraprasit et al. (2016) dataset. These authors attributed several bone elements to *Bubalus*, but we were generally unable to confirm these identifications. In part, this is because some of the fossils (e.g., DMR-KS-05-03-26-2) exceed the size of the largest modern *Bubalus* individuals in our dataset. As a consequence, the measurements frequently pointed toward an attribution to our largest taxon, *Bos gaurus*. Ratios could sometimes mitigate this effect, but as discussed above, most ratios show extensive overlap between *Bos* and *Bubalus*, and the gracility differences between these taxa are not sufficiently clear in our dataset. As a result, identifications based solely on measurements tended to be inconclusive or to reassign the remains to *B. gaurus*. For Cervidae, our results demonstrate that the identification of postcranial remains is likely far more ambiguous than previously suggested, owing to substantial overlap between the fossil measurements and those of multiple modern taxa. In several cases, however, certain elements showed strong correspondence with taxa not previously reported in the assemblage of Khok Sung, such as *Cervus nippon* and *Pseudois nayaur*. We are not able to evaluate the significance of these putative identifications of taxa not previously reported in the fossil record of Thailand, as they would need to be supported by diagnostic morphological evidence. However, they may open new avenues for future discussion of the Middle Pleistocene fauna of Thailand and show how useful the anatomical criteria developed in the next papers of this series (Chantasri et al., 2026; Mithong et al., 2026) will be. The possibility of morphological changes between modern and past representatives of the species included in our dataset should also be considered. This may affect the reliability of identification attempts, particularly when based on very limited morphometric information, and should therefore be treated with great caution.

### … and what are its limits

Our approach relies on a direct comparison between fossil and modern assemblages. For this comparison to be meaningful, the composition of both assemblages must be identical, as the method cannot easily accommodate fossil specimens belonging to taxa absent from the modern reference sample or taxa having undergone morphological changes. This limitation is clearly illustrated by our comparison with the work of Suraprasit et al. (2016), as Middle Pleistocene representatives of modern taxa may exhibit subtle morphological differences that make direct comparisons partly irrelevant. Our dataset is also not applicable to post-Neolithic assemblages because *Bos taurus* and *Bos frontalis* were not included. More generally, the method is sensitive to size differences between modern and fossil representatives of the same species. This is an important concern, as size changes have been documented in Quaternary ruminants from Southeast Asia (Hooijer, 1958), and some lineages—such as *Muntiacus*—are known to have included giant forms (Stimpson et al., 2019).

Finally, our results on limb proportions show that a simple 2D morphometric approach applied to isolated elements performs poorly compared with the same approach applied to multiple elements from the same individual. Although the method proposed in this paper is well-suited to archaeological and recent paleontological assemblages—where bones are rarely found articulated—it is not the most appropriate basic morphometric tool for identifying exceptionally well-preserved, complete specimens. In addition, our investigation was restricted to a basic comparative approach, permitting flexible input of one or several measurements from a single bone, and we did not attempt a true multivariate analysis, which would be more challenging to implement in a field study.

## Conclusion

To conclude, we present here a useful and simple approach for obtaining tentative identifications of postcranial remains of Ruminantia in Southeast Asia. Nonetheless, the method has clear limitations, as outlined above. This is not surprising: reducing the morphology of a bone to a small set of measurements is a strong simplification of its shape, and therefore restricts our ability to distinguish among taxa. The only way to overcome this limitation is to conduct more detailed morphometric analyses (e.g., 3D geometric morphometrics) or to combine measurements with qualitative morphological criteria. The latter would undoubtedly make our current approach far more effective and precise.

The osteometric dataset presented here—designed for Late Pleistocene and Holocene wild species from Southeast Asia—could be expanded with domestic species and additional wild taxa to widen its applicability across regions and time periods. However, for prehistoric studies in continental Southeast Asia, the most meaningful improvement will come from integrating this dataset with qualitative taxonomic characters. This would allow researchers to consider size, proportions, and discrete morphological features together, enabling more precise identifications and more nuanced discussions of potential size changes through time. These aspects are developed in the following papers of this series, where we present the morphological characters of the taxa included in our dataset. In these two studies, the large-sized taxa (Chantasri et al., 2026) and the small and medium-sized taxa (Mithong et al., 2026) were treated separately, following the size groups identified in the present study (Figure 1). Being of intermediate size, *Rusa unicolor* was included in both. *Tragulus* sp., the smallest taxon in the dataset, is covered by neither study as being unambiguously identified by size alone. The osteometric tool developed here can be used to strengthen attributions based on morphological criteria, to cross-check morphological and metric data, or to propose putative attributions for bones too poorly preserved to be identified on morphological grounds.

## Acknowledgements

The authors are grateful to the museums, universities, and staff members who supported this study. We especially thank the Muséum national d’Histoire naturelle (Paris, France) and Joséphine Lesur; the National Museum of Natural History (Washington, D.C., USA) and Darrin Lunde; the Zoologische Staatssammlung München (Munich, Germany) and Anneke H. van Heteren; the Museum für Naturkunde (Berlin, Germany) and Christiane Funk; and the Royal Belgian Institute of Natural Sciences (Brussels, Belgium) and Olivier Pauwels.

## Funding

This work has been done thanks to the support of two funding agencies: the Agence Nationale de la Recherche Project “ARCHAIST” (ANR n°ANR-24-CE27-5465) (dir. C. Bochaton) and the Tremplin MUSE project “ARCHECO” funded by the University of Montpellier (dir. C. Bochaton). This project was also supported by the *Mission Préhistorique Franco-Thaïe* of the French Ministry of Europe and Foreign Affairs (MEAE, Paris).

## Conflict of interest disclosure

The authors declare that they comply with the PCI rule of having no financial conflicts of interest in relation to the content of the article.

## Data, scripts, code, and supplementary information availability

Supplementary information is available online: https://doi.org/10.23708/PD4UQ6; Seesod et al., 2026.

## Author contributions

**Conceptualization**: Corentin Bochaton; **Data curation**: Corentin Bochaton; **Formal analysis**: Corentin Bochaton and Papavee Seesod; **Funding acquisition**: Corentin Bochaton; **Data acquisition**: all the authors; **Supervision**: Corentin Bochaton; **Writing original draft**: Corentin Bochaton; **Writing – review & editing**: All the authors.

## Notes

https://doi.org/10.23708/PD4UQ6

### Competing Interest Statement

The authors have declared no competing interest.

### Summary of Updates

Corrections of mistakes in the initial dataset, addition of additional comparisons with published assemblages in the text, and updates of the HTML application.

